# Individualized Gray Matter Deviations in Children with ADHD: Insights from Structural MRI Modeling

**DOI:** 10.64898/2026.03.16.710218

**Authors:** Areesha Farid, Munir Muhammad

**Affiliations:** Go Agile Cloud, Greenford, England, United Kingdom

**Keywords:** ADHD, gray matter volume, structural MRI, normative modeling, individualized deviations, prefrontal cortex, striatum, cerebellum

## Abstract

**Background:** Attention-Deficit/Hyperactivity Disorder (ADHD) affects approximately 7.6% of children globally and exhibits heterogeneous cognitive and behavioral manifestations. Conventional group-level MRI analyses often obscure individual variability in brain structure, limiting understanding of personalized neuroanatomical profiles.

**Objective:** This study quantified individualized gray matter volume (GMV) deviations in children with ADHD using age- and sex-matched normative structural MRI references.

**Methods:** Structural MRI data from 31 children with ADHD (16 males, 15 females; ages 7–15) and 413 typically developing controls (TDC; ages 7–22) were analyzed. Voxel-based morphometry extracted regional GMV across prefrontal cortex, striatal nuclei, and cerebellar vermis. Individual deviations were calculated as z-scores relative to normative distributions and categorized as typical, mild, moderate, strong, and extreme.

**Results:** Lateral and orbital prefrontal regions exhibited the highest deviations: for females, the Lateral Orbital Gyrus (LOrG) showed 33.3% mild-to-strong deviations and 13.3% extreme deviations, while the Opercular Inferior Frontal Gyrus (OpIFG) had 73.3% mild-to-strong deviations. In males, the LOrG showed 31.2% moderate, 6.2% strong, and 18.8% extreme deviations. Striatal nuclei exhibited mixed patterns: female caudate volumes were typical in 33.3% of participants, moderate-to-extreme deviations occurred in 46.7%; male putamen was typical in 31.2%, with 37.5% showing strong or extreme deviations. Cerebellar vermis values were mostly typical (50–60%) with occasional mild-to-strong deviations. Medial and superior frontal regions remained largely typical (40–73%).

**Conclusion:** Children with ADHD display heterogeneous and region-specific GMV deviations, most pronounced in lateral and orbital prefrontal cortex and select striatal regions. Individualized z-score profiling captures variability obscured in group averages, supporting personalized neuroanatomical assessment for understanding ADHD and guiding targeted treatment.

## Introduction

Attention-Deficit/Hyperactivity Disorder (ADHD) is one of the most common neurodevelopmental disorders, affecting approximately 7.6% of children aged 3–12 years globally and substantial proportions of adolescents [1]. Accurate clinical diagnosis is challenging due to overlapping symptoms with other psychiatric conditions, reliance on subjective behavioral reports, and variability in clinician practices, often leading to underdiagnosis or misdiagnosis in pediatric populations [2][3]. These challenges highlight the need for objective, standardized biomarkers to support early and reliable identification of ADHD.

Structural magnetic resonance imaging (sMRI) has been widely used to study gray matter volume (GMV) differences in ADHD. Inattentive and hyperactive/impulsive traits are linked to smaller GMV in the bilateral caudal anterior cingulate and caudate in children and adolescents [4]. Multivariate GMV network analyses show ADHD-related patterns in the putamen, caudate, accumbens, and frontal cortex that relate to attention problems [5]. Subtype studies find different GMV changes in frontal and basal ganglia regions between combined and inattentive ADHD types [6]. Comorbid oppositional defiant disorder (ODD) also affects GMV, with reduced anterior cerebellar GMV compared to ADHD alone [7]. Large studies from ENIGMA and ABCD report smaller basal ganglia (accumbens, amygdala, caudate, putamen) and hippocampal volumes in children with ADHD compared to controls [8]. These GMV differences highlight structural brain changes in ADHD and suggest MRI could help identify and understand ADHD.

Current approaches in MRI research often rely on group-level comparisons, which may not capture the full range of individual differences in brain development. Normative modeling is a statistical approach that builds reference charts of typical brain structure across age and sex and then measures how each individual differs from these norms. Wolfers et al. used normative modeling to show that individuals with ADHD often have unique patterns of deviation from typical brain structure that are not captured by group averages, highlighting the variability in brain anatomy between people with ADHD [9]. A recent large study applied normative growth models of cortical thickness to more than 1,000 children and found distinct ADHD subtypes with different structural patterns [10]. Normative brain charts have also been used to link individual differences in specific brain regions, such as the amygdala, with ADHD risk and symptom severity in adolescents [11]. Advanced normative modeling approaches have shown that individualized deviations from expected brain measures can improve understanding of how brain development differs in clinical groups compared to typical populations [12].

Thus, this study aimed to systematically quantify regional gray matter deviations in children with ADHD by comparing each individual’s brain measures to age- and sex-matched normative references. By doing so, we sought to identify subject-specific structural deviations, characterize the heterogeneity of neuroanatomical patterns in ADHD, and explore their potential relevance for personalized diagnosis and risk stratification.

## Material and Method

### Participants and Data Acquisition

This study used structural MRI data from the ADHD-200 dataset, a publicly available neuroimaging collection provided by the Neuroimaging Informatics Tools and Resources Clearinghouse (NITRC) and originally organized for the ADHD-200 Global Competition. The normative reference cohort consisted of 413 typically developing children (TDC) collected from five independent institutes, all of whom were neurologically healthy at the time of scanning. The age range of the normative cohort was 7–22 years, with representation of both sexes (194 females, 219 males). 31 children diagnosed with ADHD (16 males, 15 females; age range 7-15 years) were selected from clinical records across the same institutes for demonstration of individualized deviation analysis. The study focused on subject-level assessment, without group-level comparisons.

### MRI Protocol

All structural MRI scans were acquired on 3 Tesla Siemens scanners using 3D T1-weighted MPRAGE sequences in the sagittal plane. Acquisition parameters were restricted to maintain cross-site consistency: repetition time (TR) 2100–2530 ms, echo time (TE) 2.9–3.6 ms, inversion time (TI) 900–1200 ms, flip angle 7–10°, slice thickness 1 mm, 128–176 slices per slab. Field-of-view (FoV) was 256 mm with phase encoding anterior–posterior. Scanner and protocol consistency were maintained across sites, only scans acquired with identical hardware and acquisition protocols were retained. No retrospective harmonization of multisite data was applied.

### Inclusion and Exclusion Criteria

For the TDC cohort, inclusion criteria were:

i. No documented neurological or psychiatric diagnosis.
ii. Availability of a high-quality T1-weighted structural MRI scan.
iii. Complete demographic information including age and sex.

Participants were excluded if MRI scans exhibited visible artifacts or poor image quality, if demographic data were missing, or if scans were acquired using non-Siemens scanners or acquisition protocols inconsistent with the standard 3D T1-weighted MPRAGE sequence.

For the ADHD demonstration cohort, inclusion required a prior clinical diagnosis of ADHD and availability of a high-quality T1-weighted MRI scan. No restrictions were imposed based on ADHD subtype. Only scans meeting the same acquisition criteria as the normative cohort were retained. Because acquisition parameters were consistent across sites, no retrospective harmonization was applied.

### MRI Preprocessing and Gray Matter Volume Extraction

Structural MRI scans were preprocessed using a standardized voxel-based morphometry (VBM) pipeline implemented through DeepMRIPrep, a deep-learning framework designed to prepare T1-weighted MRI data for volumetric analysis by performing automated preprocessing steps efficiently. The pipeline included orientation normalization, bias field correction to correct for scanner-dependent intensity inhomogeneities, skull stripping to remove non-brain tissue, and tissue segmentation. Processed outputs were saved in labeled directories for reproducibility with only essential derivatives retained. These included regional volume CSV files, labeled NIfTI images of gray matter, and summary files containing total intracranial volume (TIV) and global gray matter volume (GMV). Intermediate and auxiliary files were removed to reduce storage requirements and maintain standardized outputs. For each participant, gray matter volumes were normalized by TIV to account for differences in head size.

### Brain Region Definition and Biomarker Extraction

Regional GMV was extracted using the Neuromorphometrics atlas which provides anatomically defined cortical and subcortical regions. Regions of interest (ROIs) were selected based on prior neurodevelopmental and ADHD literature, including the prefrontal cortex (middle frontal gyrus, superior frontal gyrus, orbital frontal regions, and anterior cingulate gyrus), the basal ganglia (caudate, putamen, pallidum, and nucleus accumbens), and the cerebellum (vermal lobules VIII–X). For each subject, raw regional GMV values (in mm^3^) were normalized by total intracranial volume to account for inter-individual differences in head size:

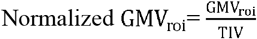

These normalized values formed the basis for subject-level deviation analysis.

### Normative Reference Modeling

To establish age- and sex-specific normative distributions, TIV-normalized GMV values of the TDC cohort were stratified by biological sex and one-year age bins. For each region, the mean and standard deviation were computed within the matched age– sex group. These statistics formed the normative reference against which individual ADHD subjects were compared. This approach provides a quantitative baseline reflecting typical developmental variability in brain structure.

### Subject-Level Deviation Analysis

Individual deviation analysis was performed for each ADHD subject by comparing their regional TIV-normalized GMV to the corresponding age–sex matched normative reference. A z-score was calculated for each region using the formula:

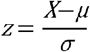

where X represents the subject’s normalized GMV, μ the mean, and σ the standard deviation of the matched normative group. Positive z-scores indicate larger GMV relative to peers, while negative z-scores indicate smaller GMV.

For higher-level interpretation, ROIs were grouped into functionally related networks, including prefrontal cortex, frontostriatal regulatory network, and cerebellar vermis. Mean z-scores were calculated within each network to summarize the overall deviation pattern.

### Interpretation of GMV Deviations

Z-scores indicate how far an individual’s gray matter volume (GMV) is from the mean of the typically developing control group, measured in standard deviations. In this study, deviations were interpreted using established thresholds: values within ±1 standard deviation of the mean were considered within the typical range, z-scores between one and ±1.5 indicated mild divergence from peers, scores between ±1.5 and ±2 represented moderate divergence, values between ±2 and ±3 indicated strong divergence, and z-scores equal to or greater than ±3 were classified as extreme divergence. These cutoffs are based on the statistical properties of the normative distribution and provide an objective framework for quantifying individual variation without implying clinical diagnosis.

## Results

Individual-level z-scores were calculated for prefrontal cortical subregions (MFG, MSFG, SFG, FRP, ACgG, OrIFG, LOrG, MOrG, OpIFG), subcortical striatal regions (Caudate, Putamen, Pallidum, Accumbens), and cerebellar vermis lobules VIII–X. Female participants were concentrated in early childhood, tween 7–10 years. Male participants showed a broader distribution between 8–12 years.

### Individual-Level Prefrontal Cortex GMV Patterns

Subject-wise z-score profiling revealed variability in prefrontal cortex (PFC) morphology among participants with ADHD.

#### 1. Female Particpants

Among the 15 female participants, regional analysis of prefrontal cortex volumes revealed heterogeneous divergence patterns. The Middle Frontal Gyrus (MFG) showed 46.7% typical values, with 26.7% mild and 13.3% extreme deviations. The Anterior Cingulate Gyrus (ACgG) remained largely typical (73.3%), though 13.3% demonstrated extreme divergence. Orbital and lateral regions exhibited comparatively higher deviation frequencies. The Lateral Orbital Gyrus (LOrG) showed 20.0% strong and 13.3% extreme deviations, while the Orbital Inferior Frontal Gyrus (OrIFG) demonstrated 33.3% mild and 13.3% strong divergence. The Opercular Inferior Frontal Gyrus (OpIFG) displayed the lowest proportion of typical values (26.7%) and a balanced distribution of mild to strong deviations (73.3% combined). Medial and superior frontal regions, including the Medial Superior Frontal Gyrus (MSFG), Superior Frontal Gyrus (SFG), and Frontal Pole (FRP), were predominantly within the typical range (40.0–53.3%), with limited extreme findings (0–6.7%).

#### 2. Male Particpants

Among the 16 male participants, the middle frontal gyrus (MFG), nearly half of the participants had values within the typical range, while 12.5% showed mild or moderate divergence, 18.8% exhibited strong deviations, and 12.5% had extreme deviations. The medial superior frontal gyrus (MSFG) displayed a higher proportion of mild divergence (37.5%), with typical values in 37.5% of participants and moderate deviations in 18.8% while strong or extreme deviations were minimal. Superior frontal gyrus (SFG) volumes were mostly typical (56.2%), but there were notable deviations, with 12.5% mild, 6.2% moderate, 12.5% strong, and 12.5% extreme. The frontal pole (FRP) was predominantly within the typical range (50.0%), though 25.0% showed mild, 12.5% moderate, and 12.5% strong divergence. Similarly, the anterior cingulate gyrus (ACgG) had 56.2% typical values, with 12.5% mild, 6.2% moderate, 18.8% strong, and 6.2% extreme deviations. Orbital and lateral regions demonstrated more frequent and pronounced deviations. The orbital inferior frontal gyrus (OrIFG) had only 37.5% typical values, with 18.8% mild, 31.2% moderate, and 12.5% strong deviations. The lateral orbital gyrus (LOrG) exhibited 25.0% typical, 18.8% mild, 31.2% moderate, 6.2% strong, and 18.8% extreme deviations, suggesting that this region is particularly prone to variation among male participants. The medial orbital gyrus (MOrG) showed a similar pattern, with only 37.5% typical, 6.2% mild, 12.5% moderate, 18.8% strong, and 25.0% extreme deviations. Finally, the opercular inferior frontal gyrus (OpIFG) presented predominantly typical values (56.2%), but 31.2% showed mild and 12.5% strong deviations, while no participants exhibited moderate or extreme divergence.

**Figure 1.**
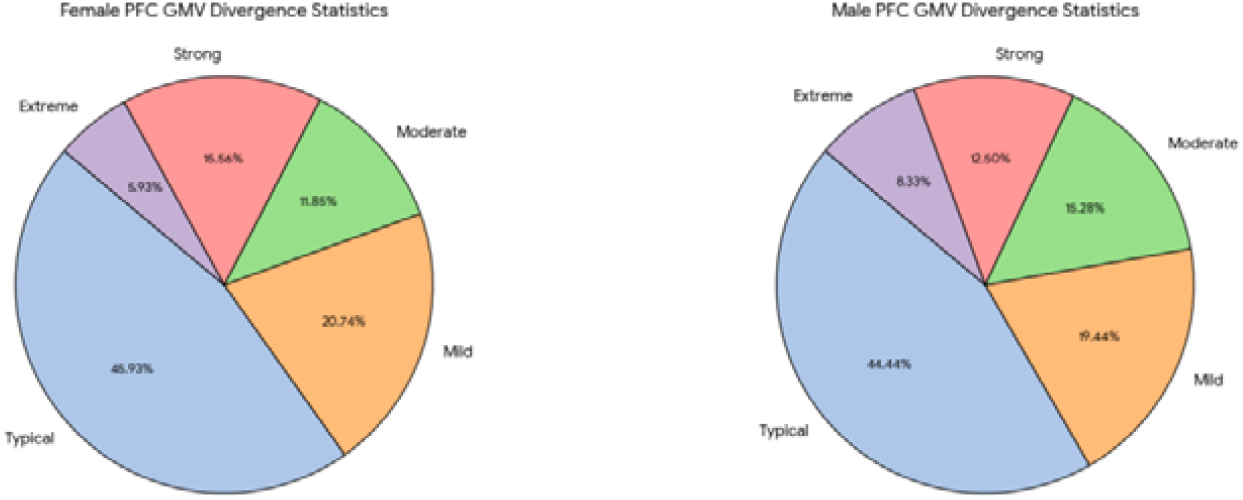
Comparison of Prefrontal Cortex (PFC) Gray Matter Volume (GMV) divergence levels across female (left) and male (right) participants with ADHD.

**Figure 1A.**
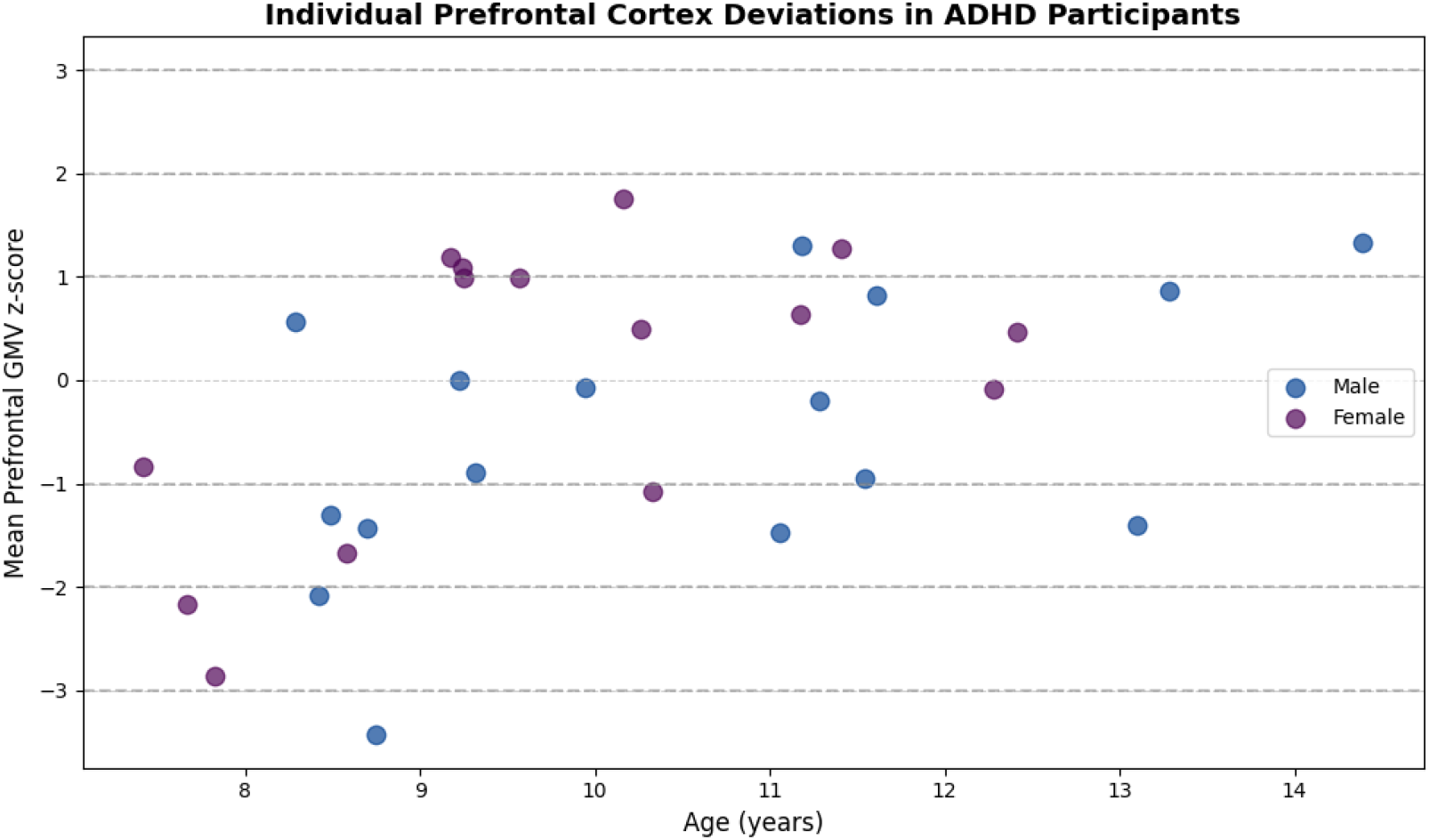
Scatter plot of participant age versus mean prefrontal GMV z-score, highlighting variability and sex distribution.

### Striatal GMV Deviations

Striatal nuclei exhibited distinct patterns compared with cortical regions, with deviations observed in both directions.

#### 1. Female Participants

In the striatum, the caudate showed mostly typical values (33.3%), with mild (20.0%), moderate (26.7%), strong (13.3%), and one extreme deviation (6.7%). The putamen displayed a more variable pattern: 33.3% typical, 13.3% mild, 20.0% moderate, and 33.3% strong deviations. The pallidum was largely typical (60.0%), with smaller proportions of mild (6.7%), moderate (13.3%), strong (13.3%), and extreme (6.7%) deviations. The accumbens exhibited 33.3% typical, 26.7% mild, 6.7% moderate, and 33.3% strong deviations.

#### 2. Male Participants

In the male striatal pattern, the caudate was predominantly typical (43.8%), with mild (18.8%), moderate (18.8%), strong (6.2%), and extreme (12.5%) deviations. The putamen showed a mixed pattern: 31.2% typical, 31.2% mild, 6.2% moderate, 25.0% strong, and 6.2% extreme. The pallidum had few typical values (12.5%), with a larger proportion of mild (43.8%), moderate (25.0%), strong (12.5%), and extreme (6.2%) deviations. The accumbens was largely typical (50.0%), with smaller contributions from mild (6.2%), moderate (12.5%), strong (12.5%), and extreme (18.8%) deviations.

**Table 1.** Sex-Stratified Striatal Gray Matter Volume Deviations in ADHD Participants.

| Sex | Region | Typical | Mild | Moderate | Strong | Extreme |
| --- | --- | --- | --- | --- | --- | --- |
| Female | Caudate | 5 | 3 | 4 | 2 | 1 |
|  | Putamen | 5 | 2 | 3 | 5 | 0 |
|  | Pallidum | 9 | 1 | 2 | 2 | 1 |
|  | Accumbens | 5 | 4 | 1 | 5 | 0 |
| Male | Caudate | 7 | 3 | 3 | 1 | 2 |
|  | Putamen | 5 | 5 | 1 | 4 | 1 |
|  | Pallidum | 2 | 7 | 4 | 2 | 1 |
|  | Accumbens | 8 | 1 | 2 | 2 | 3 |

**Figure 2.**
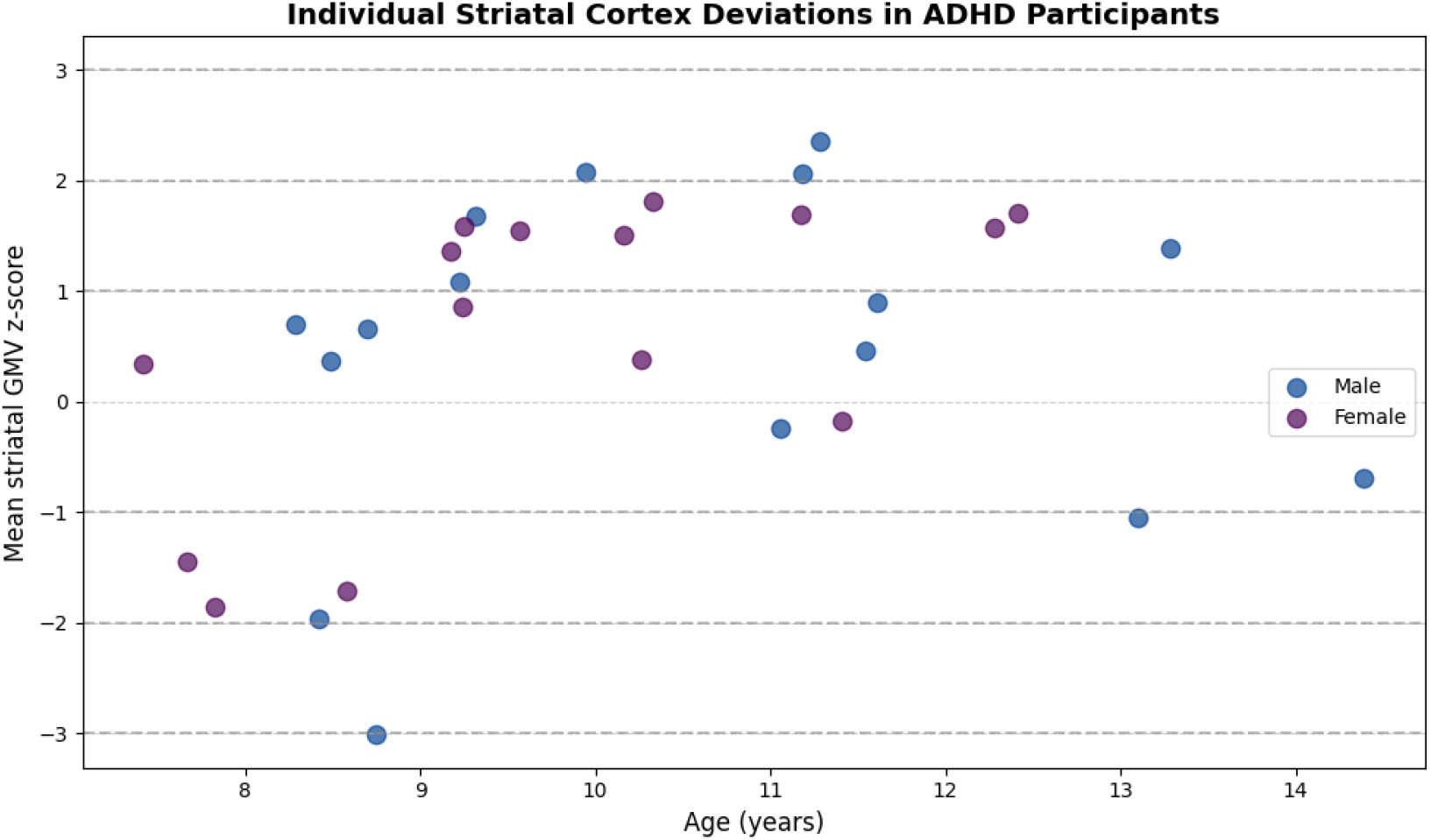
Scatter plot of age versus individual striatal GMV z-scores (Caudate, Putamen, Pallidum, Accumbens), illustrating participant-level variability and sex differences.

### Cerebellar Vermis

In the cerebellar vermis, male participants showed predominantly typical values (50.0%), with 25.0% mild, 6.2% moderate, and 18.8% strong deviations; no extreme values were observed. Among female participants, 60.0% of measurements were typical, with mild (6.7%), moderate (13.3%), and strong (20.0%) deviations, while extreme deviations were absent.

### Individualized Neuroanatomical Heterogeneity

Participant-level analysis revealed variability across prefrontal, striatal, and cerebellar regions. Medial and superior frontal areas (MSFG, SFG, FRP, ACgG) were largely typical (40–73%), while lateral and orbital prefrontal regions (LOrG, MOrG, OrIFG, OpIFG) showed the highest deviations, including mild, moderate, strong, and occasional extreme values. Striatal regions exhibited mixed patterns: the caudate and putamen showed both typical and moderate-to-strong deviations, while pallidum and accumbens had a broader range of variability. Cerebellar vermis values were mostly typical, with mild-to-strong deviations.

## Discussion

Our results revealed heterogeneous deviations in prefrontal cortical subregions, particularly in lateral and orbital areas, alongside moderate-to-strong variability in striatal regions. This pattern aligns with prior findings by Greven et al. [13], who reported reduced total gray matter and smaller caudate and putamen volumes in ADHD participants, with unaffected siblings exhibiting intermediate values. The similarity between our results and these prior observations suggests that the caudate and putamen serve as sensitive markers of ADHD and may reflect familial risk factors. Medial and superior frontal regions appeared largely typical in our cohort, whereas deviations in the lateral prefrontal cortex (PFC) were prominent. Duan et al. [14] identified gray matter networks in frontal and insular regions associated with attention and working memory deficits across adolescence and adulthood. Our findings resonate with this evidence, indicating that structural deviations in the lateral PFC may underlie persistent inattention and working memory difficulties commonly observed in ADHD. Additionally, our participant-level analyses of striatal and orbital frontal deviations align with Chen et al. [15], who reported decreased gray matter volume in the bilateral orbitofrontal cortex and right putamen among ADHD individuals.

The mixed striatal deviations observed in our cohort, particularly in the caudate and putamen, further reflect the developmental sensitivity of subcortical structures in ADHD. Hoogman et al. [16] documented smaller volumes in the caudate, putamen, accumbens, and hippocampus in children with ADHD, with differences diminishing in adulthood, consistent with our finding of variable striatal deviations across participants. Similarly, Nakao et al. [17] reported reductions in the right lentiform nucleus and caudate, moderated by age and stimulant treatment. Our participant-level z-scores suggest that striatal variability may be age-dependent and could normalize over development, supporting the concept of catch-up growth in subcortical regions. We also observed distinct patterns in striatal and cerebellar regions, with mixed deviations across participants, echoing Chang et al. [18], who reported longitudinal changes in the right striatum and cerebellar lobule VI, linking higher baseline volumes to elevated ADHD symptoms at follow-up. Furthermore, subtype-specific patterns are evident in our cohort: Saad et al. [19] demonstrated that ADHD-Inattentive (ADHD-I) individuals exhibit higher nodal degree in hippocampus and frontal regions, whereas ADHD-Combined (ADHD-C) participants show cerebellar and putamen alterations. These observations are consistent with our findings that lateral PFC and striatal regions display pronounced variability, potentially reflecting subtype-specific neuroanatomical profiles. Finally, our heterogeneous prefrontal deviations alongside variable striatal volumes correspond with Xie et al. [20], who reported smaller anterior cingulate cortex (ACC) and larger caudate in ADHD adults, contrasting with orbitofrontal and amygdala reductions in BD-I patients. The concordance of our orbital and striatal deviations with ADHD-specific alterations reinforces the importance of frontal-striatal circuits in the disorder while highlighting differential patterns relative to other neuropsychiatric conditions. The 2026 study on medication-naive offspring of parents with bipolar disorder [21] highlighted reductions in gray matter volume and cortical thickness in prefrontal and limbic regions, particularly linked to hyperactivity/impulsivity severity. This aligns with our findings, where lateral and orbital prefrontal subregions (LOrG, OrIFG, MOrG, OpIFG) showed the highest deviations among both male and female participants. The study supports the idea that ADHD-related structural variability is region-specific and modulated by genetic or developmental risk factors, reinforcing the significance of individualized z-score profiling to capture participant-level heterogeneity in cortical and subcortical networks.

Our participant-level z-score analysis revealed that lateral and orbital prefrontal regions (LOrG, MOrG, OrIFG, OpIFG) and striatal nuclei (caudate, putamen, accumbens) showed the highest deviations, while medial and superior frontal areas (MSFG, SFG, FRP, ACgG) were largely typical. These findings align with Pan et al [22], who identified ADHD structural biotypes using morphometric similarity networks, showing region-specific deviations corresponding to symptom profiles. The similarity between our z-score patterns and their network-based alterations highlights that individualized structural profiling effectively captures neuroanatomical heterogeneity underlying distinct ADHD symptom dimensions. Consistent with the precision neurodiversity framework [23], our individualized z-score analysis revealed substantial variability in prefrontal, striatal, and cerebellar GMV among ADHD participants, highlighting the importance of personalized brain network profiles over group averages. Lateral and orbital prefrontal regions (LOrG, OrIFG) showed frequent mild-to-extreme deviations, while striatal nuclei exhibited mixed patterns, reflecting individual differences in executive and cognitive networks. Cerebellar vermis values were mostly typical, suggesting some regions maintain relative normative structure despite variability. Our individualized z-score profiling of prefrontal, striatal, and cerebellar GMV deviations aligns with Pecci-Terroba et al [24], who used normative MRI centiles to identify heterogeneous ADHD subgroups. Both studies highlight that neuroanatomical alterations are region-specific and multidirectional, with lateral and orbital prefrontal regions showing the highest deviations. While Pecci-Terroba focused on subgroup centiles, our approach captures individual-level variability, emphasizing that ADHD structural patterns vary across participants.

These findings indicate that structural differences in frontal-striatal and cerebellar regions may contribute to the cognitive and behavioral features of ADHD, highlighting potential neural targets for future research and intervention. They provide further evidence that ADHD is associated with region-specific neuroanatomical variations rather than uniform brain alterations, which may help explain variability in symptoms and developmental outcomes

### Limitations

This study has several limitations. The sample size was relatively small, especially when considering age and sex subgroups, which may limit the detection of subtle brain differences. The cross-sectional design prevents conclusions about developmental trajectories or causality, and variability in ADHD presentation, including symptom severity, comorbidities, and subtype differences, could have influenced the results. Additional factors such as previous medication use, environmental influences, and MRI measurement limitations may also have affected volumetric estimates.

## Conclusion

In summary, our study demonstrates that ADHD is associated with heterogeneous structural deviations in prefrontal, striatal, and cerebellar regions, with lateral and orbital prefrontal areas showing the greatest variability. These findings support the role of frontal-striatal-cerebellar circuits in ADHD pathophysiology and highlight the importance of considering individual-level variability in neuroimaging studies. Understanding these patterns may ultimately help in predicting clinical outcomes and guiding targeted interventions.

## References

1. Salari, N., Ghasemi, H., Abdoli, N., Rahmani, A., Shiri, M. H., Hashemian, A. H., Akbari, H., & Mohammadi, M. (2023). The global prevalence of ADHD in children and adolescents: a systematic review and meta-analysis. Italian journal of pediatrics, 49(1), 48. 10.1186/s13052-023-01456-1

2. Al-Beltagi, M., Mani, B. S., Hantash, E. M., Al Zahrani, A. A., & Toema, O. (2025). Challenges in diagnosing attention-deficit/hyperactivity disorder in pediatric practice: A regional and global perspective. World journal of clinical pediatrics, 14(4), 111684. 10.5409/wjcp.v14.i4.111684

3. Sadek, J. (2023). Attention Deficit Hyperactivity Disorder Misdiagnosis: Why Medical Evaluation Should Be a Part of ADHD Assessment. Brain Sciences, 13(11), 1522. 10.3390/brainsci13111522

4. Reimann, G. E., Jeong, H. J., Durham, E. L., Archer, C., Moore, T. M., Berhe, F., Dupont, R. M., & Kaczkurkin, A. N. (2024). Gray matter volume associations in youth with ADHD features of inattention and hyperactivity/impulsivity. Human brain mapping, 45(5), e26589. 10.1002/hbm.26589

5. Wang, X. K., Wang, X. Q., Yang, X., & Yuan, L. X. (2022). Gray Matter Network Associated With Attention in Children With Attention Deficit Hyperactivity Disorder. Frontiers in psychiatry, 13, 922720. 10.3389/fpsyt.2022.922720

6. Chen, Q. R., Wang, Y., Yang, B. R., Wang, Y. F., & Chan, R. C. K. (2025). Abnormalities of gray matter volume and structural covariance in children with attention-deficit/hyperactivity disorder subtypes: implications for clinical correlations. European archives of psychiatry and clinical neuroscience, 275(7), 1897–1911. 10.1007/s00406-025-02029-5

7. Wang, X., Guo, Y., Xu, J., Xiao, Y., & Fu, Y. (2024). Decreased gray matter volume in the anterior cerebellar of attention deficit/hyperactivity disorder comorbid oppositional defiant disorder children with associated cerebellar-cerebral hyperconnectivity: insights from a combined structural MRI and resting-state fMRI study. International journal of developmental neuroscience: the official journal of the International Society for Developmental Neuroscience, 84(6), 500–509. 10.1002/jdn.10349

8. Wang, C., Wang, S., Sun, L., & Sui, J. (2026). Abnormal MRI Features in Children with ADHD: A Narrative Review of Large-Scale Studies. Brain Sciences, 16(1), 104. 10.3390/brainsci16010104

9. Wolfers, T., Beckmann, C. F., Hoogman, M., Buitelaar, J. K., Franke, B., & Marquand, A. F. (2020). Individual differences v. the average patient: mapping the heterogeneity in ADHD using normative models. Psychological medicine, 50(2), 314–323. 10.1017/S0033291719000084

10. Bu, X., Zhao, Y., Zheng, X., Fu, Z., Zhang, K., Sun, X., … & He, Y. (2024). Normative growth modeling of brain morphology reveals neuroanatomical heterogeneity and biological subtypes in children with ADHD. bioRxiv, 2024–03.

11. Nárai, Á., Hermann, P., Rádosi, A., Vakli, P., Weiss, B., Réthelyi, J. M., Bunford, N., & Vidnyánszky, Z. (2024). Amygdala Volume is Associated with ADHD Risk and Severity Beyond Comorbidities in Adolescents: Clinical Testing of Brain Chart Reference Standards. Research on child and adolescent psychopathology, 52(7), 1063–1074. 10.1007/s10802-024-01190-0

12. Mendes, S. L., Pinaya, W. H. L., Pan, P. M., Gadelha, A., Belangero, S., Jackowski, A. P., Rohde, L. A., Miguel, E. C., & Sato, J. R. (2024). GPT-based normative models of brain sMRI correlate with dimensional psychopathology. Imaging neuroscience (Cambridge, Mass.), 2, imag-2-00204. 10.1162/imag_a_00204

13. Greven, C. U., Bralten, J., Mennes, M., O’Dwyer, L., van Hulzen, K. J., Rommelse, N., Schweren, L. J., Hoekstra, P. J., Hartman, C. A., Heslenfeld, D., Oosterlaan, J., Faraone, S. V., Franke, B., Zwiers, M. P., Arias-Vasquez, A., & Buitelaar, J. K. (2015). Developmentally stable whole-brain volume reductions and developmentally sensitive caudate and putamen volume alterations in those with attention-deficit/hyperactivity disorder and their unaffected siblings. JAMA psychiatry, 72(5), 490–499. 10.1001/jamapsychiatry.2014.3162

14. Duan, K., Jiang, W., Rootes-Murdy, K., Schoenmacker, G. H., Arias-Vasquez, A., Buitelaar, J. K., Hoogman, M., Oosterlaan, J., Hoekstra, P. J., Heslenfeld, D. J., Hartman, C. A., Calhoun, V. D., Turner, J. A., & Liu, J. (2021). Gray matter networks associated with attention and working memory deficit in ADHD across adolescence and adulthood. Translational psychiatry, 11(1), 184. 10.1038/s41398-021-01301-1

15. Chen, C., Sun, S., Chen, R., Guo, Z., Tang, X., Chen, G., Chen, P., Tang, G., Huang, L., & Wang, Y. (2025). A multimodal neuroimaging meta-analysis of functional and structural brain abnormalities in attention-deficit/hyperactivity disorder. Progress in neuro-psychopharmacology & biological psychiatry, 136, 111199. 10.1016/j.pnpbp.2024.111199

16. Hoogman, M., Bralten, J., Hibar, D. P., Mennes, M., Zwiers, M. P., Schweren, L. S. J., van Hulzen, K. J. E., Medland, S. E., Shumskaya, E., Jahanshad, N., Zeeuw, P., Szekely, E., Sudre, G., Wolfers, T., Onnink, A. M. H., Dammers, J. T., Mostert, J. C., Vives-Gilabert, Y., Kohls, G., Oberwelland, E., … Franke, B. (2017). Subcortical brain volume differences in participants with attention deficit hyperactivity disorder in children and adults: a cross-sectional mega-analysis. The lancet. Psychiatry, 4(4), 310–319. 10.1016/S2215-0366(17)30049-4

17. Nakao, T., Radua, J., Rubia, K., & Mataix-Cols, D. (2011). Gray matter volume abnormalities in ADHD: voxel-based meta-analysis exploring the effects of age and stimulant medication. The American journal of psychiatry, 168(11), 1154–1163. 10.1176/appi.ajp.2011.11020281

18. Chang, J. C., Lin, H. Y., & Gau, S. S. (2024). Distinct developmental changes in regional gray matter volume and covariance in individuals with attention-deficit hyperactivity disorder: A longitudinal voxel-based morphometry study. Asian journal of psychiatry, 91, 103860. 10.1016/j.ajp.2023.103860

19. Saad, J. F., Griffiths, K. R., Kohn, M. R., Clarke, S., Williams, L. M., & Korgaonkar, M. S. (2017). Regional brain network organization distinguishes the combined and inattentive subtypes of Attention Deficit Hyperactivity Disorder. NeuroImage. Clinical, 15, 383–390. 10.1016/j.nicl.2017.05.016

20. Xie, H., Cao, Y., Long, X., Xiao, H., Wang, X., Qiu, C., & Jia, Z. (2022). A comparative study of gray matter volumetric alterations in adults with attention deficit hyperactivity disorder and bipolar disorder type I. Journal of psychiatric research, 155, 410–419. 10.1016/j.jpsychires.2022.09.015

21. Lu, R., Deng, Y., Qin, Y., Lou, Y., Zeng, H., Tan, S., Xu, L., Xu, G., & Lv, Y. (2026). Alterations in gray matter volume and cortical thickness in medication-naive offspring with ADHD and a parental history of bipolar disorder. Journal of affective disorders, 393(Pt A), 120322. 10.1016/j.jad.2025.120322

22. Pan, N., Long, Y., Qin, K., Pope, I. Z., Chen, Q., Zhu, Z., Cao, Y., Li, L., Singh, M. K., McNamara, R. K., DelBello, M. P., Chen, Y., Fornito, A., & Gong, Q. (2026). Mapping ADHD Heterogeneity and Biotypes by Topological Deviations in Morphometric Similarity Networks. JAMA psychiatry, e260001. Advance online publication. 10.1001/jamapsychiatry.2026.0001

23. Mohammad SI, Azzam ER, Vasudevan A, Ismail SM, Ayaz H and Prasad KDV (2025) Precision neurodiversity: personalized brain network architecture as a window into cognitive variability. Front. Hum. Neurosci. 19:1669431. doi: 10.3389/fnhum.2025.1669431

24. Pecci-Terroba, C., Lai, M. C., Lombardo, M. V., Chakrabarti, B., Ruigrok, A. N. V., Suckling, J., Anagnostou, E., Lerch, J. P., Taylor, M. J., Nicolson, R., Georgiades, S., Crosbie, J., Schachar, R., Kelley, E., Jones, J., Arnold, P. D., Seidlitz, J., Alexander-Bloch, A. F., Bullmore, E. T., Baron-Cohen, S., … Bethlehem, R. A. I. (2025). Subgrouping autism and ADHD based on structural MRI population modelling centiles. Molecular autism, 16(1), 33. 10.1186/s13229-025-00667-z

